# In Situ, Antibody-Independent, and Multiplexed Characterization of Amyloid Plaques by MALDI MS/MS Imaging Using iprm-PASEF

**DOI:** 10.1101/2025.09.04.674173

**Authors:** Larissa Chiara Meyer, Mujia Jenny Li, Nadine Meier, Beate Vollmer-Kary, Tobias Feilen, Julie Audebert, Konrad Kurowski, Stephan Singer, Peter Bonsert, Melanie Christine Föll, Oliver Schilling

**Affiliations:** Institute for Surgical Pathology, Faculty of Medicine, University Medical Centre Freiburg, University of Freiburg, Freiburg, Germany; Faculty of Biology, University of Freiburg, Freiburg, Germany; Institute for Pharmaceutical Sciences, University of Freiburg, Freiburg, Germany; Institute of Pathology, University of Tübingen, Tübingen, Germany; Core Facility for Histopathology and Digital Pathology, Medical Center, University of Freiburg, Freiburg, Germany; German Cancer Consortium (DKTK), German Cancer Research Center (DKFZ), Heidelberg, Germany

## Abstract

**Background:** Amyloidosis collectively describes a heterogeneous group of protein aggregation-based diseases involving the misfolding and extracellular accumulation of fibril-forming amyloid proteins. The tissue deposition of these fibrils occurs either localized (limited to one organ) or systemically, impairing organ function and, in severe cases, leading to organ failure. Identification of the amyloidogenic protein leading to the correct subtyping is challenging but crucial for treatment decisions. Imaging parallel reaction monitoring (iprm)-parallel accumulation-serial fragmentation (PASEF) is a novel method for matrix-assisted laser desorption/ionization (MALDI)-mass spectrometry imaging (MSI) that enables the fast, spatially resolved, multiplexed, and antibody-independent identification of peptides and proteins in situ. In this proof-of-concept study, we demonstrate the applicability of iprm-PASEF for the *in situ* characterization of amyloidosis.

**Methods:** A multiplexed iprm-PASEF panel (comprising trapped ion mobility spectrometry (TIMS) and m/z values) for characterizing amyloid plaques by MALDI-MSI was compiled. The panel included nine peptide entries of amyloidosis-associated proteins (vitronectin, apolipoprotein E, serum amyloid P component) and the prevalent amyloidosis-subtype proteins serum amyloid A, transthyretin, and immunoglobulin light chain 2. The multiplexed iprm-PASEF panel was applied to a tissue microarray (TMA) assembled from formalin-fixed, paraffin-embedded (FFPE) specimens, comprising biopsies of amyloidosis-positive tissues from 18 patients, including six different tissue types, and representing the following amyloidosis subtypes: ATTR (transthyretin amyloidosis), AL (immunoglobulin light chain amyloidosis), and AA (secondary amyloidosis). For comparison and co-localization, congo red staining was performed on adjacent slides.

**Results:** MALDI TIMS MS1 imaging was acquired from the amyloidosis TMA, providing an *m*/*z* feature list containing 1390 entries that was subsequently refined to only include *m*/*z* features likely stemming from amyloidosis-related peptides. Further, we focused on optimal usage of the *m*/*z* and ion mobility range to increase the number of targeted peptides. The final iprm-PASEF panel comprised 10 entries ranging from *m*/*z* 887.51 - 1,986.85 and 1/*K*_0_ 1.4 - 2.1 V·s/cm^2^; targeting nine peptides from six different amyloidosis-related proteins. Apolipoprotein E, serum amyloid P component, and immunoglobulin light chain 1/2 were included with two entries. Transthyretin, vitronectin, and serum amyloid A were covered by one peptide. One entry was included as a positive control targeting actin A. We applied the iprm-PASEF (MS2 mode) panel to the amyloidosis TMA and fragment (MS2) spectra were analyzed by the search engine MASCOT. Eight out of ten peptides derived from vitronectin, apolipoprotein E, serum amyloid A, serum amyloid P component, and transthyretin were successfully identified with MASCOT scores above 18. Comparison to Congo red staining indicated a more localized and heterogeneous pattern of amyloid-associated proteins (vitronectin, apolipoprotein E, serum amyloid P component) by iprm-PASEF. Compared to the clinical annotation, transthyretin and serum amyloid A were only found in their respective subtypes. In total, it was possible to describe the spatial distribution of five different amyloid-associated proteins, thereby further characterizing Congo red-positive amyloid plaques in one single iprm-PASEF measurement.

**Conclusion and perspective:** This proof-of-concept study demonstrates feasibility for in situ, antibody-independent, and multiplexed proteomic typing of amyloidosis plaques. Future enlargement of the multiplex panel is likely to increase the number of targetable amyloid proteins and to probe for the presence of post-translational modifications. The present study highlights the value of iprm-PASEF for MALDI imaging strategies.

## 1 Introduction

Amyloidosis collectively describes a heterogeneous group of protein aggregation-based diseases involving the misfolding and extracellular accumulation of fibril-forming amyloid proteins. The tissue deposition of these fibrils occurs either localized (limited to one organ) or systemically, impairing organ function and, in severe cases, leading to organ failure. Diagnosing amyloidosis remains challenging, as evidence of over 40 amyloid-associated and amyloidogenic proteins leads to a variety of subtypes with heterogeneous symptoms, disease severity, and treatment regimens [1, 2]. The predominant subtypes of amyloidosis include transthyretin (ATTR) amyloidosis (caused by the accumulation of transthyretin monomers), immunoglobulin light chain amyloidosis (AL) (caused by misfolded and/or over-produced and/or incompletely processed immunoglobulin light chains), and the secondary AA amyloidosis (caused by the misfolding and accumulation of serum amyloid A (SAA)). In addition to these subtype-specific proteins, several proteins are found in most amyloid deposits and often referred to as amyloid-associated proteins. Examples include apolipoprotein E (APOE), serum amyloid P component (SAMP), and vitronectin (VTN) [3-7].

Amyloid plaques in tissue samples (e.g., biopsies) are initially detected by staining with Congo red, an azo dye that stains beta-sheet structures commonly observed in misfolded amyloid proteins. Following a Congo red-positive diagnosis, amyloidosis subtyping is required, involving the identification of the accumulated protein within the Congo red-positive areas, typically by means of immunohistochemistry [8] or dissection-based proteomics using liquid chromatography electrospray ionization tandem mass spectrometry (LC-ESI-MS/MS) [3, 9-11]. Both approaches face drawbacks: Immunohistochemistry requires the availability of suitable antibody panels, may consume one slide of limited sample material per staining, and may face limitations with regard to quantification [12]. On the other side, LC-ESI-MS/MS has the potential to profile the entire set of amyloidogenic proteins with minimal sample input. Yet, extensive sample preparation and ideally laser capture microdissection (LCM) is required to focus on spatially defined, Congo red-positive areas [13-15]. As a result, LC-ESI-MS/MS based characterization of amyloidosis is laborious.

Matrix-assisted laser desorption/ionization (MALDI) imaging of peptides offers a rapid, spatial, and antibody-independent method to map amyloidogenic peptides. In fact, MALDI imaging has already been applied to amyloidosis in several studies [16-19]. However, in situ and straightforward peptide sequence identification by peptide-spectrum-matching still remains challenging in MALDI imaging. To this end, the aforementioned MALDI imaging studies have used in situ MS2 measurements in a serial manner for selected precursor ions or subsequent LC-ESI-MS/MS analysis with matching of precursor ion masses. However, the first approach defeats the necessity of multiplexed probing for the presence of potential amyloidogenic proteins while the second approach leads to additional laborious tasks and unreliable peptide identifications.

Recently, we presented imaging parallel reaction monitoring (iprm-) Parallel Accumulation-Serial Fragmentation (PASEF) by utilizing iprm-PASEF-enabled, multiplexed, and MS/MS-based MALDI imaging of peptides that preserves the spatial information on MS2 level [20]. In this method, precursor ions are selected after a classical MS1 imaging scan of the tissue of interest and targeted in a subsequent iprm-PASEF survey, thereby providing the MS2 spectra of the precursor at every single pixel. The iprm-PASEF method exploits TIMS to separate precursors not only by their *m*/*z* value but also by their 1/*K*_0_ thus multiplexing and measuring them in one single MS/MS MALDI imaging run.

In the present proof-of-concept study, we designed a multiplexed and amyloidosis specific iprm-PASEF panel targeting nine peptides from five amyloidogenic proteins. This panel was applied to a tissue microarray (TMA) containing three different amyloidosis subtypes derived from a total of 18 patients, thus presenting the first application of iprm-PASEF for the in situ characterization of amyloidosis plaques.

## 2 Material and Methods

### 2.1 Sample collection

The Ethics Committee of the University Medical Center Freiburg approved the sample collection and usage for the amyloidosis TMA (Ref.: 20/1119). The TMA includes 34 Congo red-positive biopsies (2 mm diameter) from a total of 18 patients with clinically diagnosed amyloidosis subtypes. All patients gave written informed consent for the use of their tissues for research purposes. Before inclusion, all patient data were pseudonymized. The biopsies were collected from different tissue types including skeletal muscle, lung, salivary gland/soft tissue, liver, amyloidoma and kidney. The tissue type distribution of the TMA is shown in Figure S1. The represented amyloidosis subtypes include transthyretin (ATTR) amyloidosis, immunoglobulin light chain amyloidosis (AL; κ and λ), and the secondary AA amyloidosis (serum amyloid A (SAA)). The amyloidosis subtype was retrieved from clinical records. For two cores (NA), this information was not available.

### 2.2 Sample Preparation for MALDI Imaging of Tryptic Peptides

Sample preparation was carried out as published previously [20]. In short, two adjacent tissue slides were prepared for MALDI imaging measurement. Deparaffinization was performed using xylol and ethanol/water dilutions. The sections were washed with ammonium bicarbonate solution and underwent antigen retrieval in a citric acid buffer at ∼100°C for 1 hour in a steamer. After drying, trypsin (0.1 mg/mL in 10% ACN and 40 mM ammonium bicarbonate) was sprayed onto the slides using an M3+ sprayer (HTX Technologies, Chapel Hill, NC, USA), followed by incubation at 50 °C for 2 hours. Alpha-cyano-4-hydroxycinnamic acid matrix (10 mg/mL in 70% ACN, 7 mM ammonium phosphate and 0.1% TFA) was then applied using the same sprayer device. The spraying parameters were set according to [20].

### 2.3 MALDI TIMS MS1 Imaging Acquisition, Data Processing and Data Analysis

MALDI TIMS MS1 imaging was performed as published previously [20]. In short, MALDI TIMS MS1 imaging acquisition was performed using a timsTOF fleX mass spectrometer (Bruker Daltonics, Bremen, Germany) in positive-ion reflector mode with a 50×50 μm pixel size, 600 shots per pixel, and a 10 kHz laser frequency. Tuning parameters were as published [20]. The data was analyzed using SCiLS Lab software (Bruker Daltonics, Bremen, Germany), with an *m*/*z* interval width of 10 ppm and 0.01 V·s/cm^2^ for 1/*K*_0_. Root mean square normalization was applied. Feature finding was performed with T-ReX3 using spatial smoothing (2×2), 50% coverage, and a 1% relative intensity threshold. Manual peak selection was used when necessary.

### 2.4 Literature Search of Representative Peptide Sequences for iprm-PASEF Panel Design

As a foundation of the iprm-PASEF panel, we sought to retrieve experimentally derived *m*/*z* values for peptides of amyloidogenic proteins as determined by MALDI-MS. Literature search was performed by searching PubMed and Google Scholar in 12/2024 using the following keywords: “MALDI imaging peptide amyloidosis”, “MALDI MSI peptide imaging amyloid deposits amyloidosis”, “SAA amyloidosis MALDI imaging” and “AL amyloidosis MALDI imaging”. Ultimately, eight publications were screened for *m*/*z* values with annotated peptide identifications derived from amyloid-associated and amyloidogenic proteins [16-19, 21-24]. We focused on publications that included MALDI in situ MS/MS for peptide identification and did not consider publications without a clear description of the identification method (Table S1).

### 2.5 iprm-PASEF Acquisition, Data Processing and Data Analysis

As described previously [20], the iprm-PASEF dataset was acquired using the iprm-PASEF tool for timsControl 6.0.8, provided by Bruker Daltonics. Measurements were performed in prm-PASEF and positive-ion reflector mode on a timsTOF fleX with a 100×100 μm pixel size, 2,000 shots per pixel, and a 5 kHz laser frequency. The *m*/*z*-based isolation and fragmentations settings were chosen as prm-PASEF settings. The *m*/*z* range was 100–2,000, and the 1/*K*_0_ range was 1.2–2.1 V·s/cm^2^ with a 230 ms ramp time. Optimized tuning parameters included collision RF (750 Vpp), ion energy (5 eV), transfer time (55 ms), and pre-pulse storage (8 µs). TIMS cell pressure and 1/*K*_0_ calibration followed standard MALDI TIMS MS1 imaging procedures. The iprm-PASEF method targeted nine precursor ions with predefined 1/*K*_0_ and *m*/*z* ranges, using collision energies from 45–140 eV based on precursor mass. Thus, obtained MS2 spectra from the iprm-PASEF datasets were averaged across all pixels in DataAnalysis software and exported as “.mgf” files. These were submitted to the MASCOT MS/MS Ions Search against the Swiss-Prot database (April 2024) for mus musculus and homo sapiens. Trypsin digestion with up to one missed cleavage was allowed, with a precursor tolerance of 100 ppm, fragment ion tolerance of 0.3 Da, and a 1% FDR. Searches were performed with and without the variable modifications of oxidation of histidine, tryptophan, proline, and methionine. MS2 spectra and theoretical b- and y-ions (matched within 0.3 Da) were visualized using pyteomics in Python [25]. Peptide fragment ion images were plotted in SCiLS Lab.

### 2.6 LC-ESI-TIMS-MS/MS Sample Preparation, Acquisition and Data Analysis Focused on Singly Charged Peptides

As previously described [20] after MALDI imaging, co-crystallized peptides were extracted by dissolving the matrix layer in 70% acetonitrile, followed by vacuum drying and peptide cleanup using PreOmics Cartridges. The cleaned peptides were dissolved in water with 0.1% formic acid and desalted on Evotips before LC-ESI-TIMS-MS/MS analysis. LC-ESI-TIMS-MS/MS was performed on a timsTOF fleX mass spectrometer with a CaptiveSpray ion source, using the Evosep One HPLC system with a 44-minute gradient and a reversed-phase column. Samples were analyzed in positive data-dependent acquisition (DDA)-PASEF mode, with a mass range of 100–1,700 *m*/*z* and a 1/*K*_0_ range of 0.7–2.1 V·s/cm^2^. The TIMS cell pressure and ion mobility calibration were adjusted to enhance the detection of singly charged peptides. Key tuning parameters were optimized, including a capillary voltage of 1,600 V, drying gas flow of 3 L/min at 180°C, and collision energy of 20–75 eV.

Data analysis was performed using the FragPipe pipeline (v22.0). MS2 peptide identification was done with MSFragger, setting hydroxylation (+15.999 Da) of proline as a variable modification and using Bruker.d raw data as input. Human and mouse protein databases (including contaminants and internal standards) were used, with reversed sequences as decoys.

## 3 Results

### 3.1 iprm-PASEF of Amyloidosis TMA

#### 3.1.1 MS1 Imaging Acquisition

In this proof-of-concept study, the applicability of iprm-PASEF to identify a small panel of amyloidosis-associated peptides was tested on a TMA comprising tissue samples of different amyloidosis subtypes (ATTR, AL (κ), AL (λ), AA). MS1 tryptic peptide imaging (50×50 µm spatial resolution) of the amyloidosis TMA comprising 34 tissue cores derived from 18 patients took 230 minutes and generated 48,311 pixels, resulting in a speed of approximately 3.5 pixels per second, which is less than 200 min per cm^2^ of tissue. 1,390 features (MS1 peaks which are defined by a specific *m*/*z* value and a potential 1/*K*_0_ value) were found using the SCiLS Lab Software.

#### 3.1.2 Precursor Selection for Amyloid Peptide-Targeting iprm-PASEF Panel Design

The iprm-PASEF tool allows multiplexed and efficient MS/MS acquisition in MALDI imaging (Fig 2A). However, it is restricted to a maximum of 15 precursors in one measurement when using the *m*/*z*-based isolation and fragmentation mode. Therefore, it is necessary to shorten the list of 1,390 features identified in the MS1 measurement. To accomplish this, an iprm-PASEF panel was designed that can identify and characterize amyloidosis-diseased tissue in a single MALDI imaging measurement. To achieve this, the literature was searched to compile a list of potential *m*/*z* values for analyzing amyloid-associated and amyloidogenic proteins and their respective peptides as described in detail in section 2.4. Only publications that performed tryptic peptide MALDI imaging and employed an in situ MS/MS method for peptide identification were included. A detailed list can be found in Table S1. Each collected *m*/*z* value was further investigated in the MS1 imaging dataset by signal intensity and distinct ion image distribution. In addition, signals were examined for overshadowing by high-intensity signals (including isotopic signals) derived from structural proteins not associated with amyloidosis, such as collagen or actin. For this purpose, the ion images were compared with those of known collagen or actin peaks (e.g. *m*/*z* 836.4425 [26, 27] or *m*/*z* 1,105.5731 [26, 28] for collagen and *m*/*z* 1,198.7024 [29] for actin A). Thereby, the *m*/*z* list obtained from literature was further refined. As the multiplexed iprm-PASEF requires non-overlapping 1/*K*_0_ windows and the precursor ion entries are presently limited by the software to 15 for the *m*/*z*-based isolation and fragmentation, the number of selectable precursors is restricted. Hence, we aimed to cover amyloid-associated proteins with at least one peptide, simultaneously targeting as many different proteins as possible within the restriction by isolation window availability. Additionally, one precursor at *m*/*z* 1,198.7024 (frequently reported as actin A (ACTA) in literature [29]) was included due to availability of isolation window area as a positive control for the measurement. The resulting precursor list and selected isolation windows can be found in Figure 2 B.

**Figure 1:**
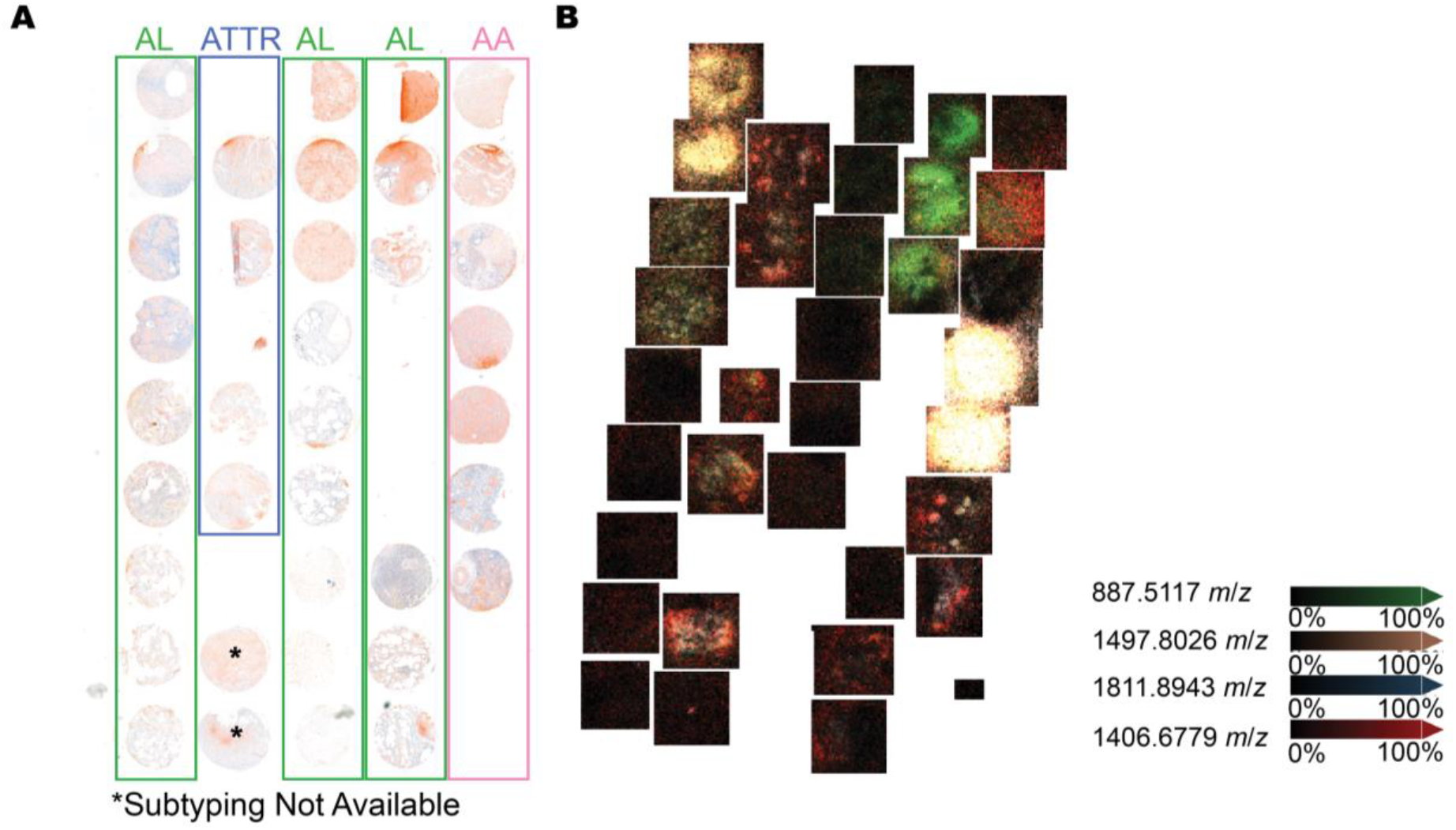
Overview of the amyloidosis TMA composition. **(A)** Congo red staining of the amyloidosis TMA. The TMA includes 34 biopsies (2 mm diameter) from a total of 18 patients. The represented amyloidosis subtypes include transthyretin (ATTR, blue) amyloidosis, immunoglobulin light chain amyloidosis (AL; κ and λ, green), and the secondary AA amyloidosis (serum amyloid A (SAA), pink). **(B)** Four exemplary selected ion images of an MS1 MALDI imaging run of the amyloidosis TMA overlayed. The squares equal the selected measurement regions containing the tissue cores.

**Figure 2:**
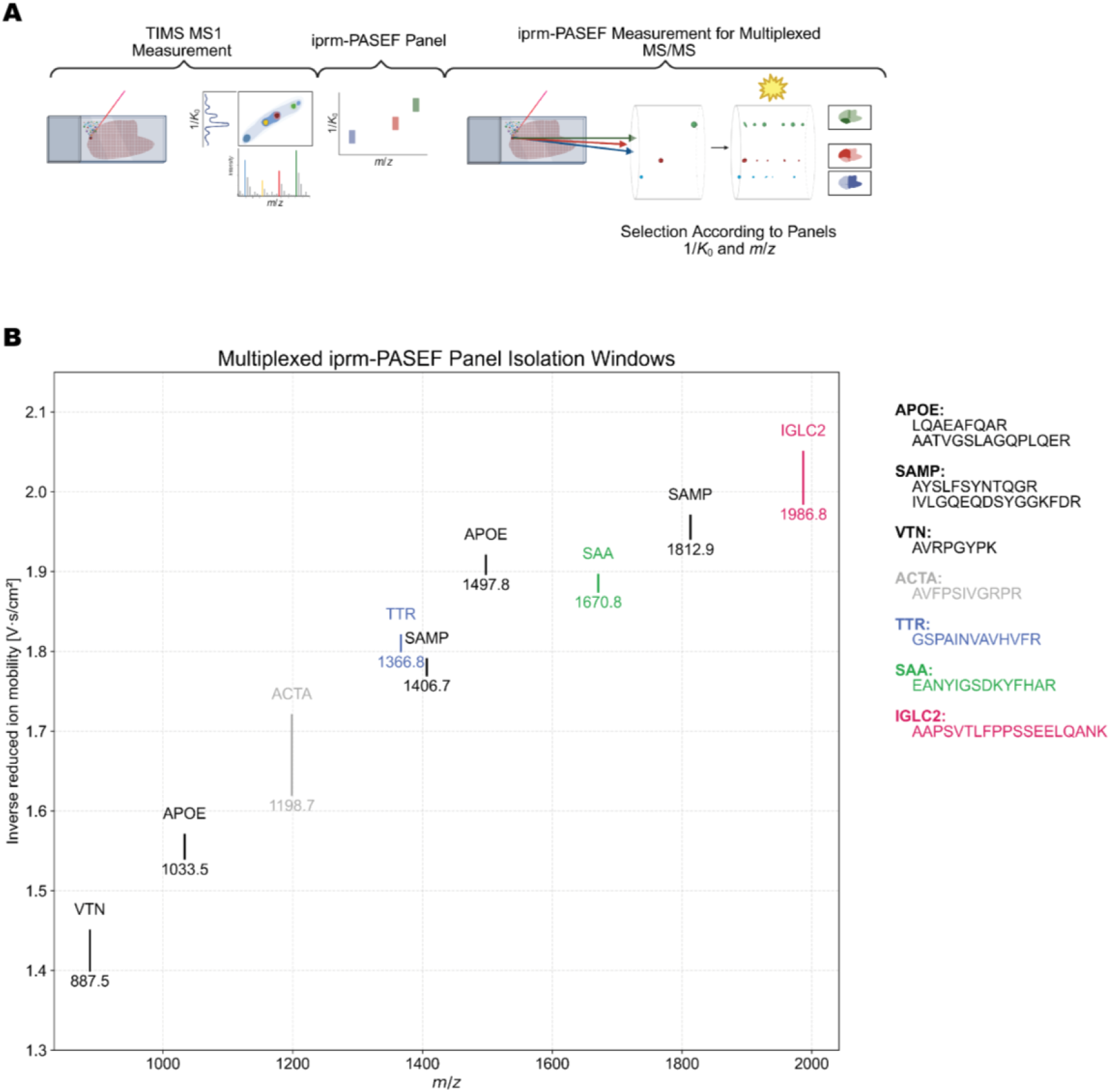
Isolation windows of selected precursors for the multiplexed iprm-PASEF panel. **(A)** Overview of the iprm-PASEF method used for this study. **(B)** The *m*/*z* and ion mobility windows for all selected precursors are visualized, highlighting the usage of the entire available ion mobility range.

We are aware that literature-based peptide annotations provide only limited confidence, mainly due to the inability of MS1 *m*/*z* values alone to distinguish between isobaric peptides. Therefore, a subsequent iprm-PASEF measurement was performed to confirm correct peptide identities by in situ MS/MS directly on the tissue of interest and after ion mobility separation.

#### 3.1.3 MS/MS Imaging By iprm-PASEF

The iprm-PASEF measurement was performed with a pixel size of 100×100 µm on an adjacent TMA slide using the aforementioned precursor list (Figure 2B). The measurement generated 13,667 pixels in 112 minutes, resulting in approximately 2 pixels per second which is less than 100 min per cm^2^ of tissue. Bruker.d raw data was imported into Data Analysis (Bruker Daltonics, Bremen, Germany) and automatically split according to precursor-specific ion-mobility traces. The resulting mean MS2 spectra were exported in a.mgf format and submitted to the online MS/MS ion search engine from MASCOT, allowing for peptide identification. In total, eight out of nine precursors from the iprm-PASEF measurement were successfully identified by MASCOT. The MASCOT score, provided by Matrix Science, statistically evaluates peptide-to-spectrum matching (PSM) confidence, with higher scores indicating greater reliability. In this study, scores ranged from 18.62 to 44.19 with a mass accuracy lower than 35 ppm, consistent with those reported in previous in situ MALDI MS/MS studies [30-33].

Interestingly, two additional features at *m/z* 886.4770 and *m/z* 886.4414 were co-isolated with the precursor at *m/z* 887.5069 (identified as VTN). The fragment signals were separated by their ion mobility trace and three distinct MS2 spectra were extracted and analyzed successfully by MASCOT (Figure S2) as peptides assigned to COL1A1 and APOE and are included in Table 1. This observation underlines the usefulness of ion mobility separation in MALDI imaging.

**Table 1.**
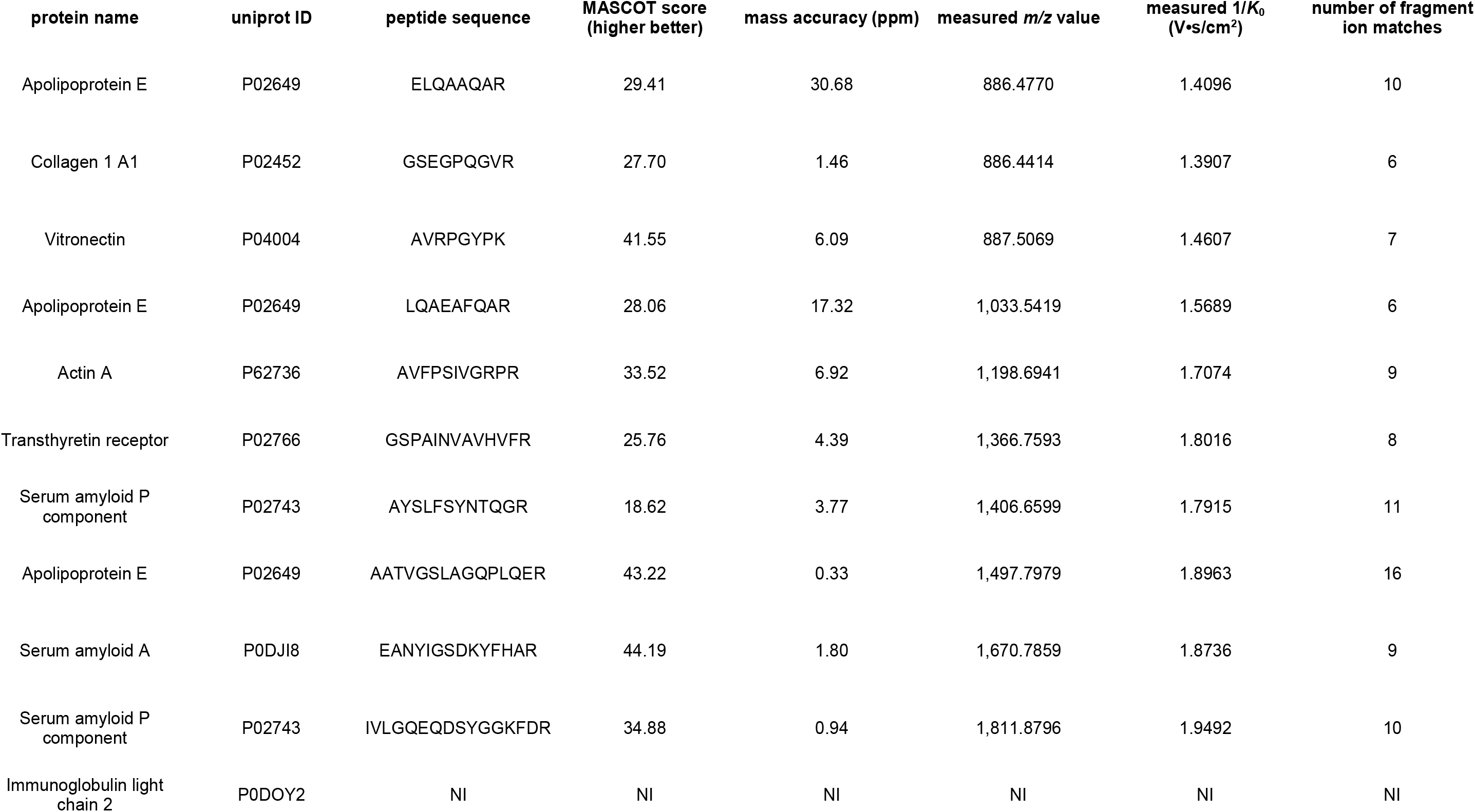
Validations of peptide identities detected in iprm-PASEF of an amyloidosis TMA. NI= not identified.

The isolation window targeting an immunoglobulin light chain-derived peptide at *m*/*z* 1,986.8565 and 2.0179 V·s/cm^2^ isolated an MS2 spectrum that includes the fragment signal at *m*/*z* 175.1186, indicating arginine as a y1-ion. However, we were not able to identify the isolated tandem mass spectra by MASCOT and the identity remains elusive.

In summary, we identified 10 peptides in one single iprm-PASEF run by using the MASCOT search engine (Table 1).

#### 3.1.4 Further Corroboration of Peptide Identities

We further sought to corroborate the peptide identities obtained by iprm-PASEF and MASCOT on the spatial level. To this end, we annotated the predicted b- and y-ions of the mean MS2 spectra using the pyteomics tool [25].

Then, the co-localization of matched b- and y-ions to the respective precursor was investigated in SCiLS Lab to assess the spatial distribution of the MS/MS information. We visually assessed the co-localization of the targeted precursors with their expected b- and y-ions in the MS2 ion images (difference of maximum 15 ppm) (Figure S4). For example, the precursor with the peptide sequence EANYIGSDKYFHAR and the *m*/*z* value of 1,670.7859 showed a similar spatial distribution to its matched fragment ions (Figure 3).

**Figure 3:**
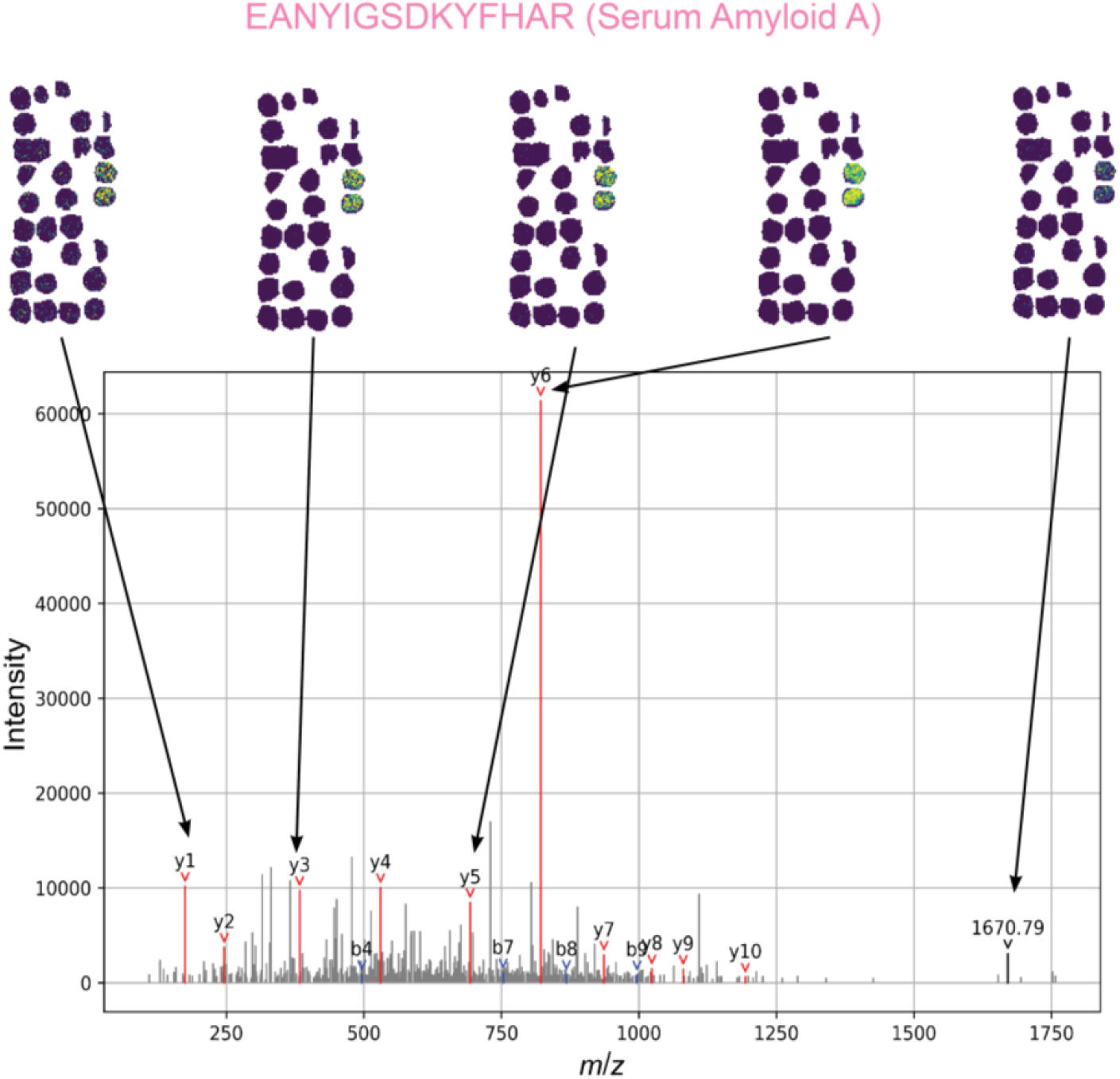
MS2 spectrum of the targeted SAA peptide EANYIGSDKYFHAR. Mean MS2 spectrum acquired in iprm-PASEF with a spatial resolution of 100 µm is shown for one selected precursor of the amyloidosis TMA. B-ions are highlighted in red and y-ions in blue. Ion images of respective fragments are shown on top of the spectrum.

To further corroborate peptide identification, residual matrix after iprm-PASEF was used to extract peptides for LC-ESI-TIMS-MS/MS measurement. The measurement method included singly charged peptides to allow for the comparison on 1/*K*_0_ ion mobility values in addition to *m*/*z* values. 9 out of 10 iprm-PASEF identified peptides could be detected in LC-ESI-MS/MS with an *m*/*z* variance of less than 10 ppm. Five of them were even detected as singly charged ions, enabling the direct matching of *m*/*z* and 1/*K*_0_ values. The observed 1/*K*_0_ variance between MALDI and ESI was lower than 0.025 V·s/cm^2^ (Table S2).

### 3.2 iprm-PASEF Allows for the In Situ, Antibody-Independent Characterization of Amyloidosis Subtypes

Having established a small-scale iprm-PASEF amyloidosis panel, we investigated to what extent MSI-based characterization of amyloidosis subtypes correlates with their clinical-pathological diagnosis. To this end, the acquired iprm-PASEF dataset was aligned with the clinical annotations extracted from the corresponding medical records. MS1 imaging and iprm-PASEF datasets were co-localized visually with a consecutive Congo red staining. For the following analysis, we only inspected MSI signals within the Congo red-positive areas of the tissue cores.

As expected, the targeted peptides from the amyloid-associated proteins APOE, SAMP and VTN were found in all subtypes (Fig 4 A). For instance, SAMP is commonly found in most amyloid cores [3, 5, 6]. In our TMA, we detected two SAMP derived peptides in high abundance in two cores, and in low abundance in 19 out of 37 cores over all subtypes.

**Figure 4:**
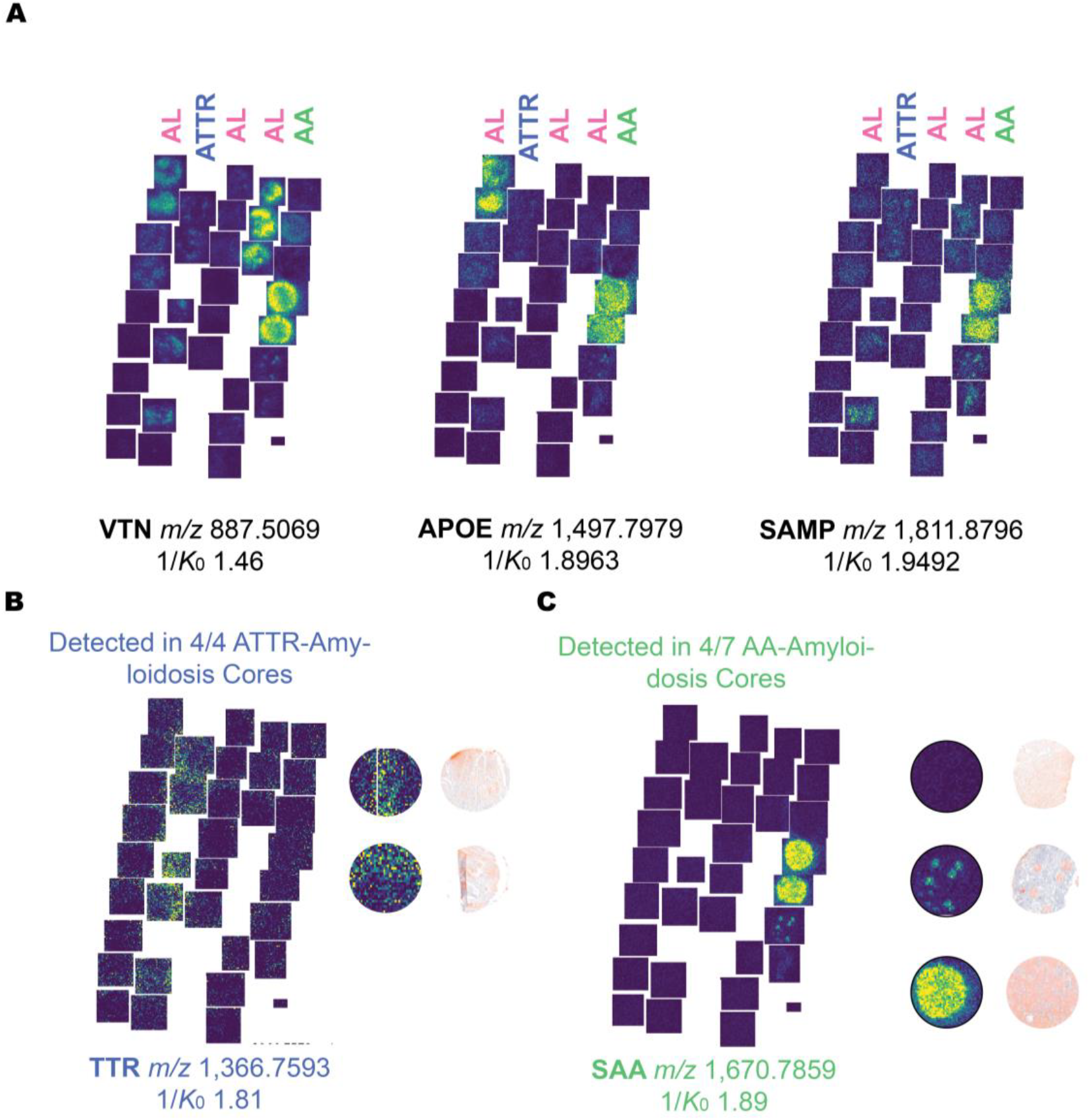
Characterization and subtyping of the amyloidosis TMA by MALDI imaging. **(A)** Characterization of amyloidosis-related proteins with targeted peptides (APOE, SAMP, VTN). **(B)** Proteins and their targeted peptides for subtyping ATTR-amyloidosis (TTHY). Congo red staining of some selected cores are visualized for comparison. **(C)** Proteins and their targeted peptides for subtyping AA-amyloidosis (SAA). Congo red staining of some selected cores are visualised for comparison.

On the other hand, the protein APOE is usually found at low levels in some AA and AL amyloidosis cores [3-5]. In this study, peptides from APOE were found in high abundance in two out of seven AA amyloidosis cores and in low abundance in four out of nine AL amyloidosis cores. Most importantly, we identified several APOE and SAMP peptides with a similar location within the tissue, confirming that the spatial resolution of the protein remains intact even after digestion into peptides.

We further targeted one peptide of the protein VTN that is reported to be present in most other amyloidosis subtypes [3]. Within our TMA, it was detected with high abundance in two out of seven AA amyloidosis cores and in three out of six AL λ cases.

Regarding the targeting of amyloidosis subtype specific peptides, we identified the peptide GSPAINVAVHVFR (residues 42-54 of TTR) by iprm-PASEF with low intensity, indicating weak expression of the protein TTR in all four ATTR-amyloidosis cores (Fig 4 B). The peptide was not detected in the cores assigned to the other subtypes AL and AA, thus showing high specificity.

Notably, this TMA contains two amyloidosis cores for which subtyping is unknown. We detected TTR expression in one of these cores, which suggests that ATTR may be the underlying subtype. Meanwhile, the subtype-specific peptide EANYIGSDKYFHAR (residues 44-57 of SAA) was identified in four out of seven AA-amyloidosis cores with varying intensity, while remaining absent in AL and ATTR cores showing high specificity for AA amyloidosis. Most interestingly, the congo red-positive area in these SAA-containing cores shows close alignment with the ion image of the SAA peptide detected by the iprm-PASEF panel with MALDI imaging (Fig 4 C). For the AL-amyloidosis subtype, the IGLC2 peptide could not be identified within the presented iprm-PASEF setup and awaits further investigation.

In total, we identified 9 peptides derived from 5 different amyloid-related proteins by applying our designed iprm-PASEF panel. Thereby, heterogenous amyloidosis-diseased tissues were spatially characterized by their composition of subtype specific and amyloid-associated proteins.

### 3.3 TIMS-Enabled Separation of Isobaric Signals

By the inclusion of the ion mobility, TIMS-included MALDI imaging measurements can lead to an increased signal resolution and enable deconvolution of features, e.g. as shown for the co-isolated peptides at m/z 887.5069 for the iprm-PASEF panel and as shown in further published studies [34, 35].

When looking closer at the MS1 signal detected at *m*/*z* 1,033.5418 in our experiment, we noticed that the *m*/*z* peak was divided into two signals in the ion mobility dimension with two different 1/*K*_0_ values (1.51 V·s/cm^2^ and 1.54 V·s/cm^2^). Interestingly, those two signals showed divergent ion images, indicating that the peak is derived from two distinct isobaric molecules and the mobility difference is not solely caused by conformational change (Figure 5). The signal at 1.54 V·s/cm^2^ was assigned to ELQAAQAR from APOE (Mascot score 29) in the iprm-PASEF measurement (3.1.3). To further investigate the identity of *m*/*z* 1,033.5418 at 1.51 V·s/cm^2^, we used the iprm-PASEF mode for MS/MS identification in an additional measurement, revealing the signal at 1.51 V·s/cm^2^ as the first isotope peak of *m*/*z* 1032.5418, detected at *m*/*z* 1,033.5418, and identified as TSGGAGGLGSLR from desmin (DES, P17661, Mascot score 45).

**Figure 5:**
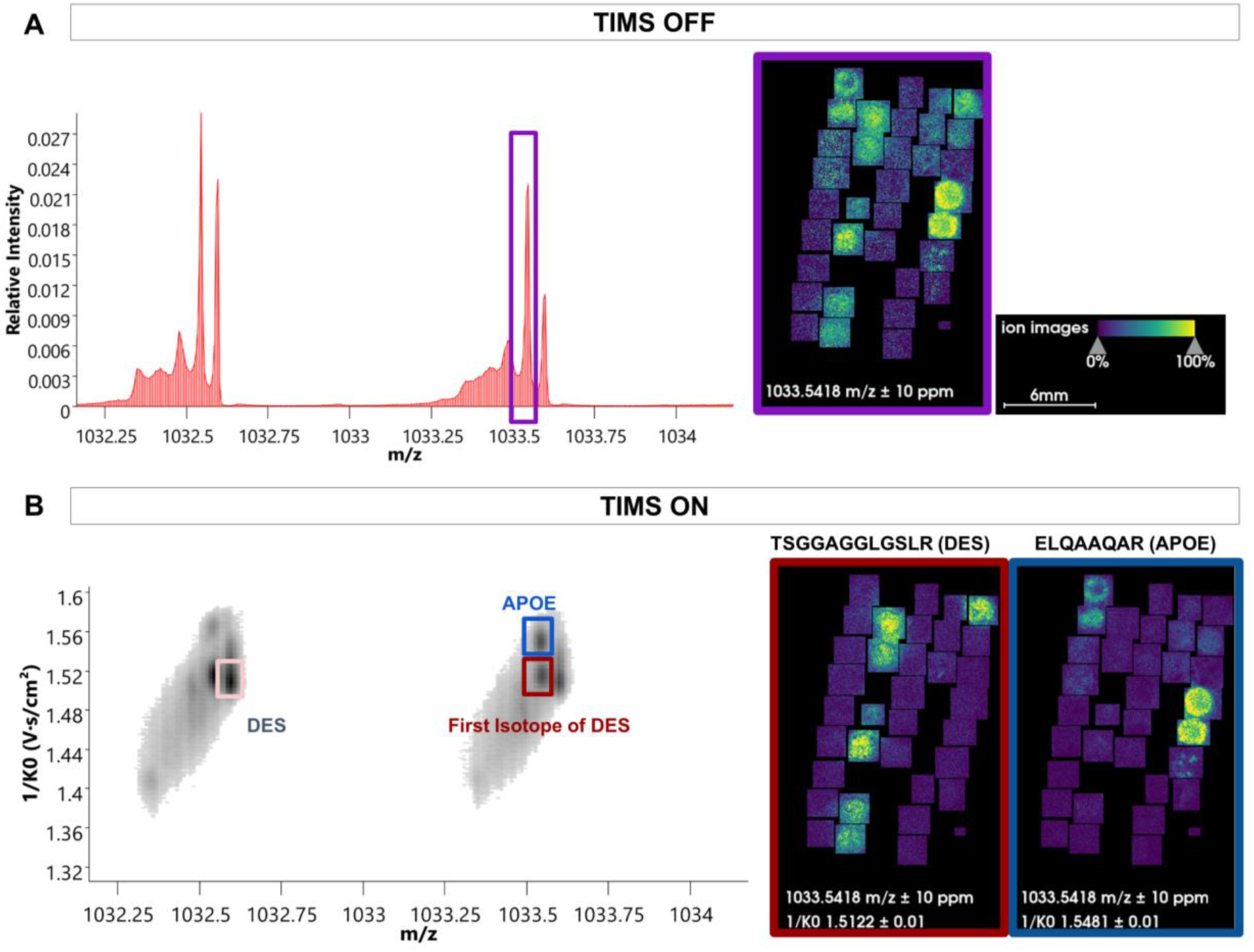
TIMS-enabled separation of isobaric signals. **(A)** Peak at *m*/*z* 1033.5 unfiltered for ion mobility shows a diffuse localisation over the whole tissue. **(B)** By including the TIMS dimension a second precursor is noticed underlying the *m*/*z* 1,033.5 peak with two different ion mobilities (1.51 V·s/cm^2^ and 1.54 V·s/cm^2^). When targeting the peaks with iprm-PASEF, APOE and the first isotope peak of DES were identified. The ion images show a much clearer localisation when TIMS dimension is included into the measurement.

This result underscores the advantage of incorporating the TIMS dimension into MALDI imaging of tryptic peptides; in the MS1 MALDI TIMS imaging, *m*/*z* 1,033.5418, referred to as APOE signal in literature, would have compiled a background signal derived by desmin, possibly leading to a false positive result for APOE in certain cores, as illustrated in the ion images in Figure 5. This also highlights the advantage of full spatial coverage for MS/MS acquisition. Overall, by incorporating the TIMS dimension, we were able to successfully isolate the APOE signal at *m*/*z* 1,033.5418 from interfering signals, thereby improving both the resolution and reliability of the peptide MALDI imaging measurements.

## 4 Discussion

In this study, we present a novel iprm-PASEF panel for characterizing the protein composition of amyloid plaques in situ. This proof-of-concept study identified a total of ten peptides derived from seven proteins, in particular five proteins related to amyloidosis disease, in a single MALDI imaging measurement and visualized their spatial distribution. Furthermore, we have demonstrated the advantage of including the TIMS dimension into MALDI imaging measurements, thereby separating isobaric signals and performing iprm-PASEF for identification.

In our iprm-PASEF panel, we targeted a total of three peptides derived from subtype specific proteins (SAA, IGLC2 and TTR). Two of those peptides were successfully identified within their respective clinical subtype (AA and ATTR), with variability observed between patients and tissue types. We manually evaluated the spatial similarities between the identified peptides and the Congo red staining. In the future, this step could be performed with a less biased approach, such as by applying an automated co-registration pipeline [36]. The significant variability observed among patients and tissue types may be explained by the varying accessibility of heat-induced antigen retrieval to different tissue structures or inefficient trypsin digestion [37]. We failed to identify the third targeted subtype-specific peptide (IGLC2), which is usually found in AL amyloidosis; this is despite its identification by additional LC-ESI-MS/MS measurements. We speculate that the underlying proteins have a too low abundance in the presented TMA to be detected and identified correctly. Further studies and possibly AL-dediciated iprm-PASEF panels may be required to yield their identification in MALDI imaging.

All five amyloidosis-associated peptides included in the iprm-PASEF panel could be successfully identified. SAMP and APOE were covered with up to two peptides per protein that all showed a high alignment in spatial resolution within one protein species. Although this approach was important for confirming the spatial localization of proteins after digestion, future approaches could replace these targets with other precursors of interest e.g. other peptides specifically derived from AL-amyloidosis. This would optimize the use of the limited multiplexing capacity per acquisition cycle (approximately ten), since maintaining non-overlapping 1/*K*_0_ isolation windows is currently still necessary. One way to address this issue and thereby increasing the number of targetable precursors per scan is to use a raster offset to schedule multiple isolation window panels. This would allow the same tissue sample to be acquired repeatedly. Alternatively, one could acquire in situ MS/MS information spotwise and automatically using the spatial ion mobility-scheduled exhaustive fragmentation (SIMSEF) tool [38]. This approach has the potential to significantly increase the number of precursors submitted for MS/MS fragmentation in a single run but comes at the cost of strongly reduced spatial MS/MS coverage.

The presented workflow can be further improved by establishing dedicated software for analyzing iprm-PASEF datasets. As outlined in our previous publication, our data analysis pipeline relied on a combination of several different tools, thereby entering multiple import and export steps that introduce inefficiencies and potential sources of error [20]. In addition, manual adjustments were necessary, for example due to the limited performance of the SCiLS Lab feature detection tool for 1/*K*_0_ dimensional data. Furthermore, this study only used the mean MS2 spectrum across the entire tissue region for peptide identification, which could potentially introduce a significant noise level. Integrating precursor-specific regions could substantially improve the quality of the MS2 spectra, resulting in more reliable identification. Additionally, MS2 spectrum quality is highly dependent on the mobilogram region selected for integration in DataAnalysis, indicating that multiple peptide species may have been co-isolated within a single window. Therefore, by automating the data analysis pipeline and using more sophisticated and adapted software e.g for PSM, identification and throughput could be increased.

In conclusion, this study applies an amyloidosis-targeting iprm-PASEF panel to a TMA derived from 18 patients, thus identifying and spatially mapping amyloidosis-associated peptides in a multiplexed manner. This allows for a more refined characterization of congo red positive amyloid plaques. It further opens novel possibilities for the usage of MALDI imaging in spatial proteomics.

## Abbreviations

ACTA: Actin A
APOE: Apolipoprotein E
FFPE: Formalin-fixed paraffin-embedded
IGLC2: Imunglobulin Light Chain 2
iprm-PASEF: Imaging Parallel Reaction Monitoring, Parallel Accumulation SErial Fragmentation
LC-ESI-MS/MS: Liquid-chromatography electrospray ionization tandem mass spectrometry
LCM: Laser Capture Microdissection
MALDI: Matrix-assisted laser desorption/ionization
MSI: Mass spectrometry imaging
prm: Parallel Reaction Monitoring
PASEF: Parallel Accumulation SErial Fragmentation
PSM: Peptide to spectrum matching
SAA: Serum Amyloid A
SAMP: Serum Amyloid P Component
TIMS: Trapped Ion Mobility Spectrometry
TMA: Tissue Microarray
VTN: Vitonectin
1/*K*_0_: Inverse of reduced ion mobility

## Supplementary Notes

**Figure S1**: Overview of the tissue types included in the amyloidosis TMA.

**Figure S2**: Identification of two distinct precursors in one isolation window: *m*/*z* 887.51 and m/z 886.47 were isolated in a single iprm-PASEF window and successfully extracted separately by their mobility separation. The signal *m*/*z* 887.51 (green) could be identified as a peptide AVRPGYPK (VTN) by MASCOT and *m*/*z* 886.47 as ELQAAQAR (APOE).

**Figure S3**: Exported mean MS2 spectra from the iprm-PASEF dataset, one for each isolation window submitted.

**Figure S4**: Ionimages of targeted precursors and their respective detected fragment ions in the MS2 iprm-PASEF measurement.

**Table S1**: Literature Search for *m*/*z* values of amyloidosis-related peptides detected in MALDI imaging

**Table S2:** Targeted peptides detected in the the subsequent LC-ESI-TIMS-MS/MS Measurement, ND = Not detected

