## Supplementary Figures for "In Situ, Antibody-Independent, and Multiplexed Characterization of Amyloid Plaques by MALDI MS/MS Imaging Using iprm-PASEF"

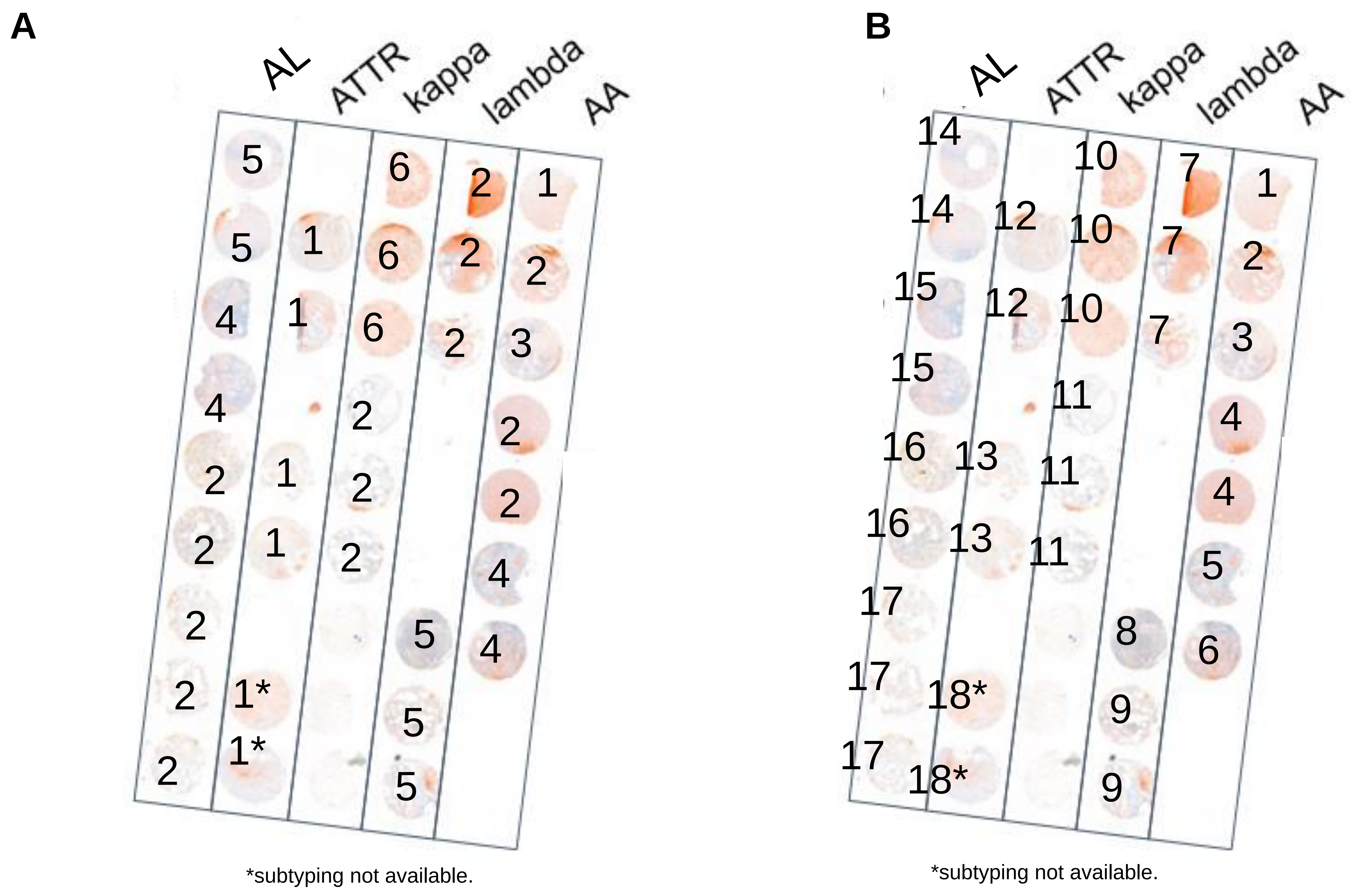


1: Sceletal muscle
2: Lung
3: Salivary gland/soft tissue
4: Kidney
5: Liver
6: Amyloidoma

Patient Numbers

**Figure S1**: Detailed annotations of the amyloidosis TMA. (A) Tissue subtypes. (B) Patient replicates.

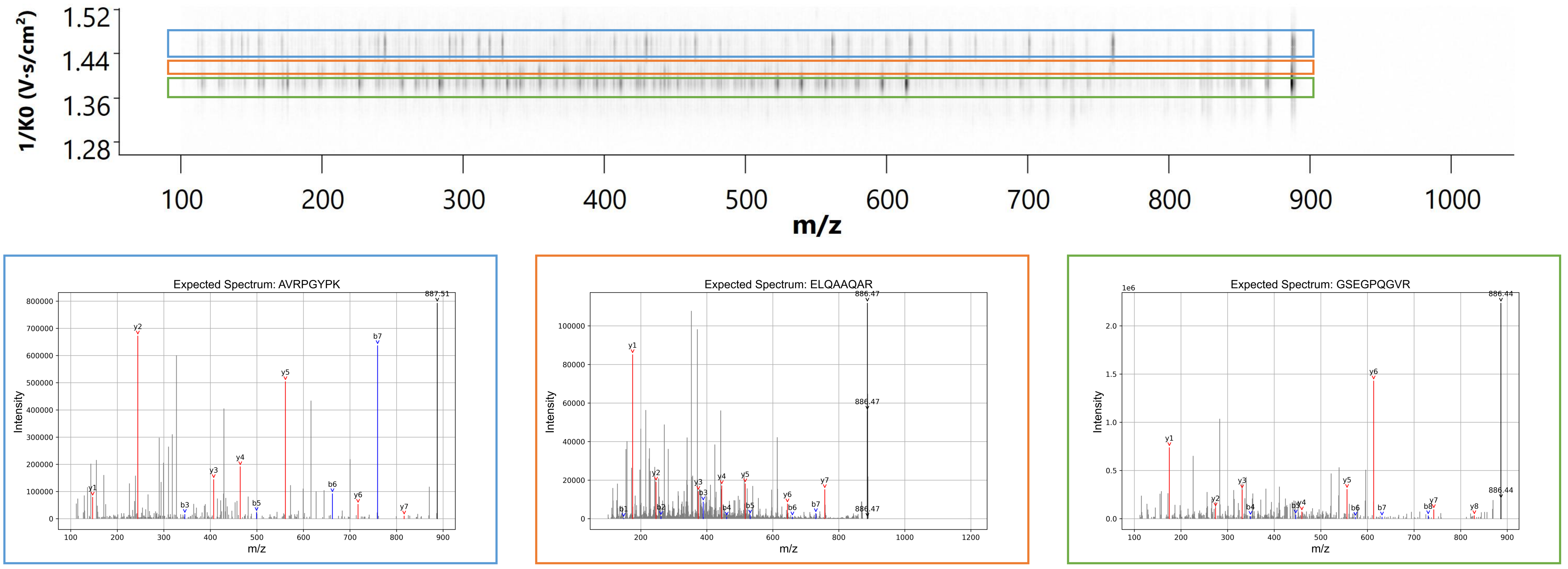


**Figure S2:** Identification of two distinct precursors in one isolation window: *m/z* 887.51 and m/z 886.47 were isolated in a single iprm-PASEF window and successfully extracted separately by their mobility separation. The signal *m/z* 887.51 (green) could be identified as a peptide AVRPGYPK (VTN) by MASCOT and *m/z* 886.47 as ELQAAQAR (APOE).

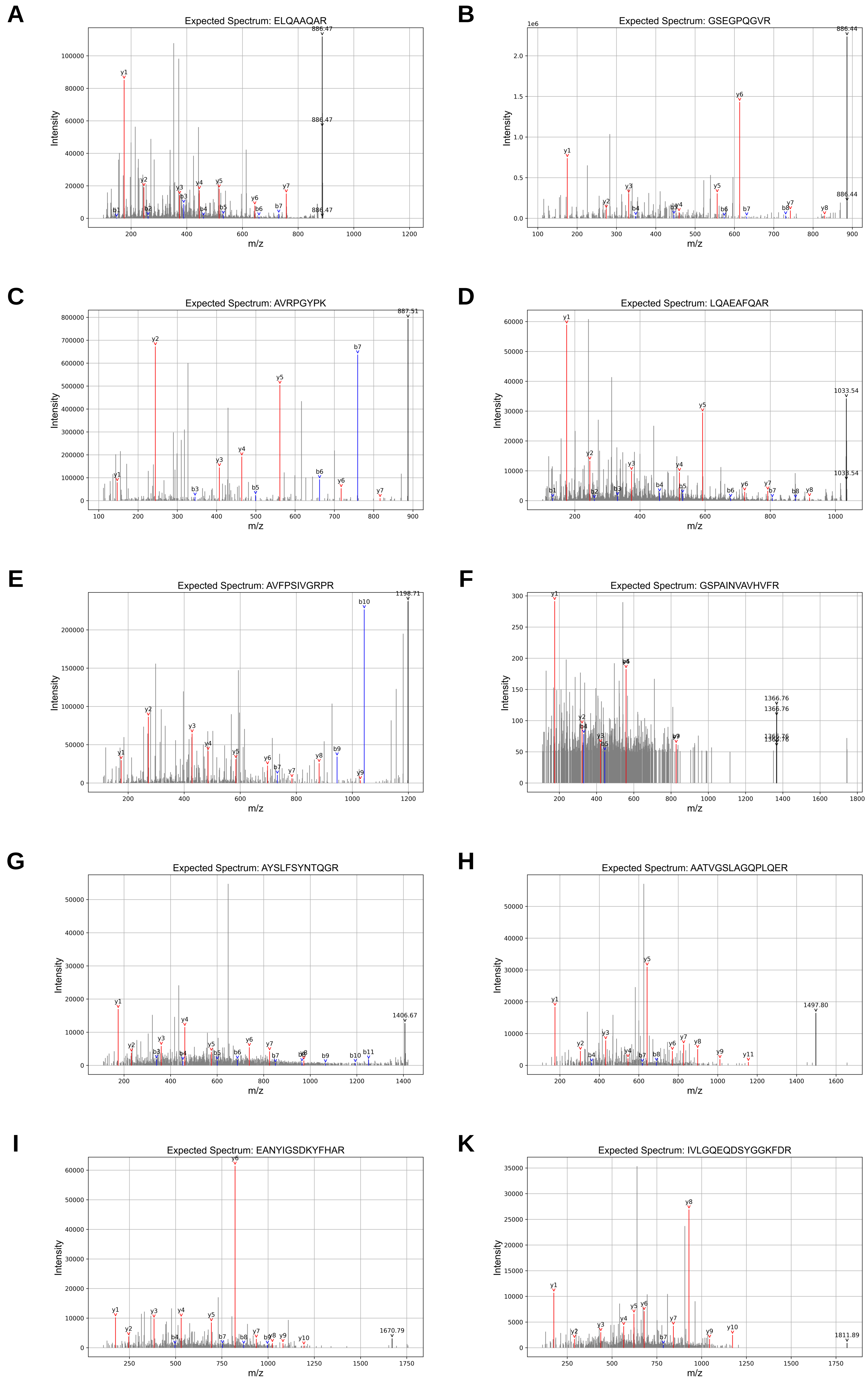


**Figure S3:** Exported mean MS2 spectra from the iprm-PASEF dataset, one for each isolation window submitted.

886 APOE

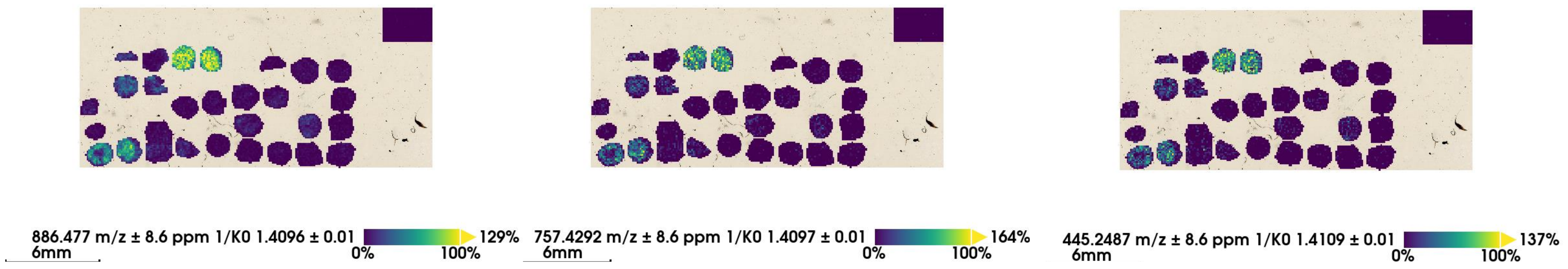


886 COL1A1

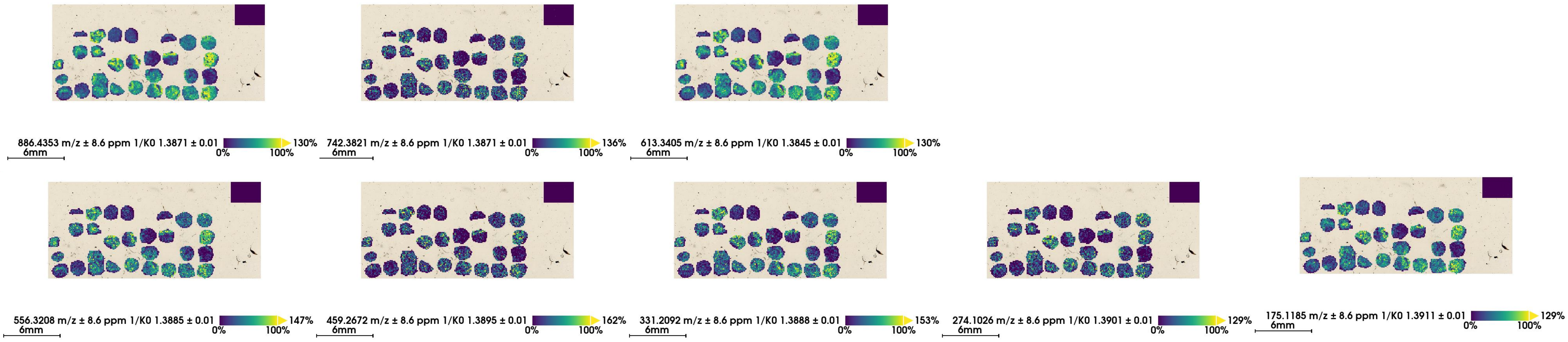


887 VTN

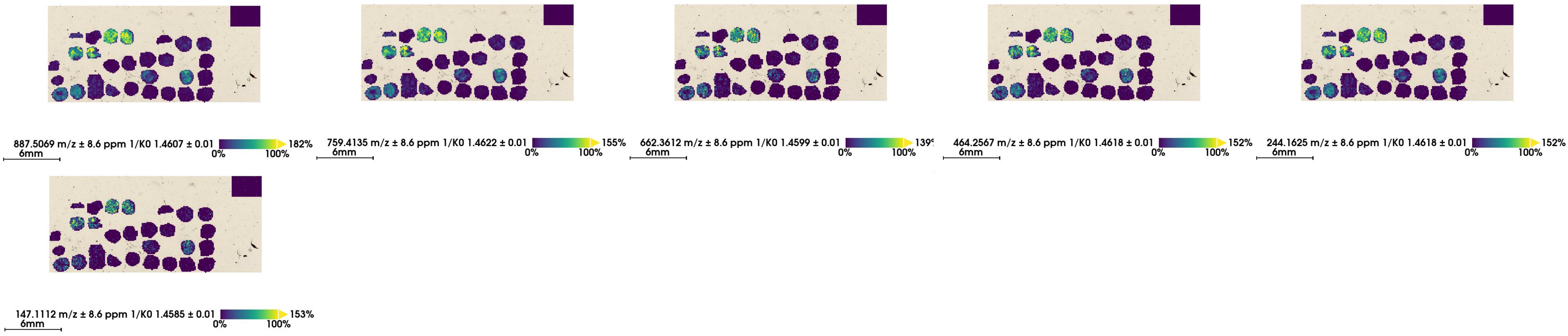


1033 APOE

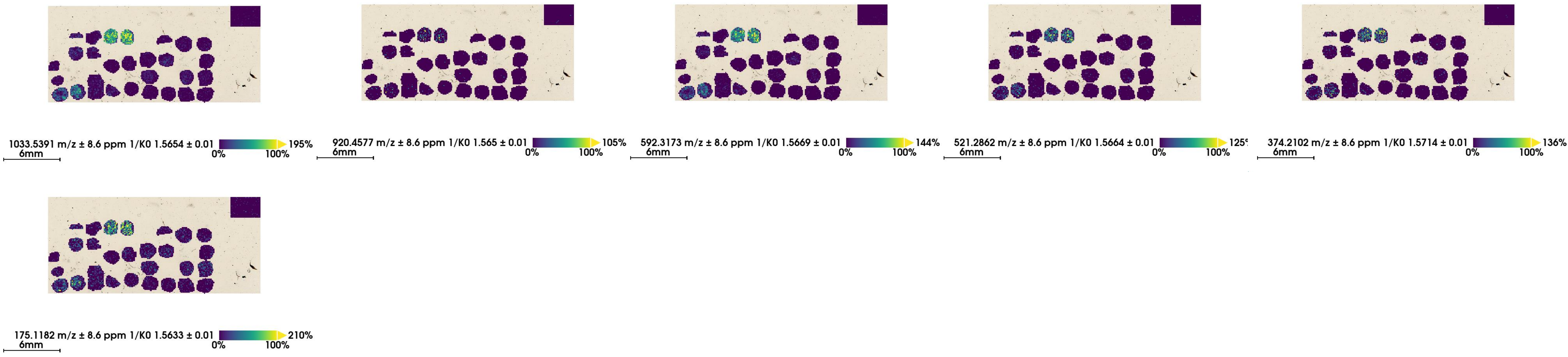


1198 ACTA

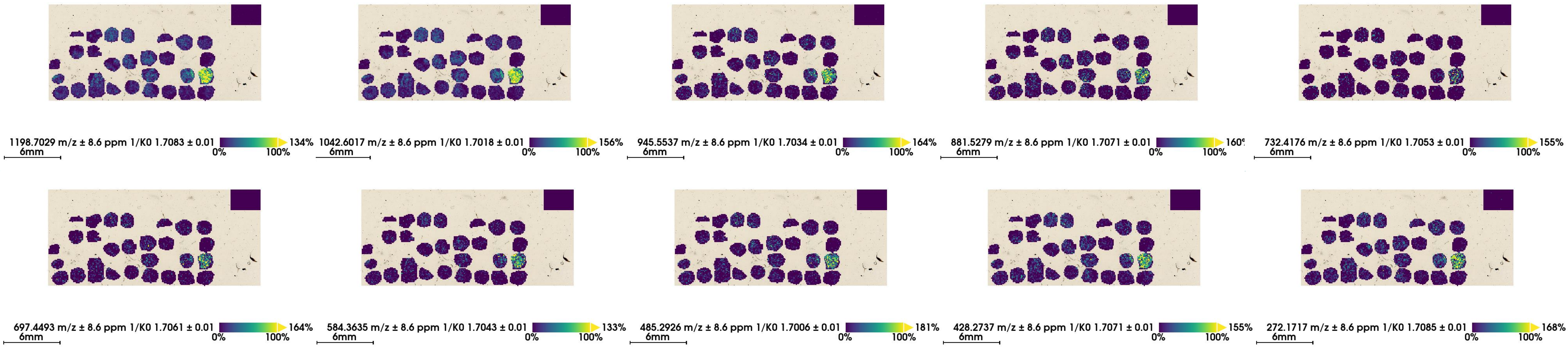


**Figure S4**: Ionimages of targeted precursors and their respective detected fragment ions in the MS2 iprm-PASEF measurement.

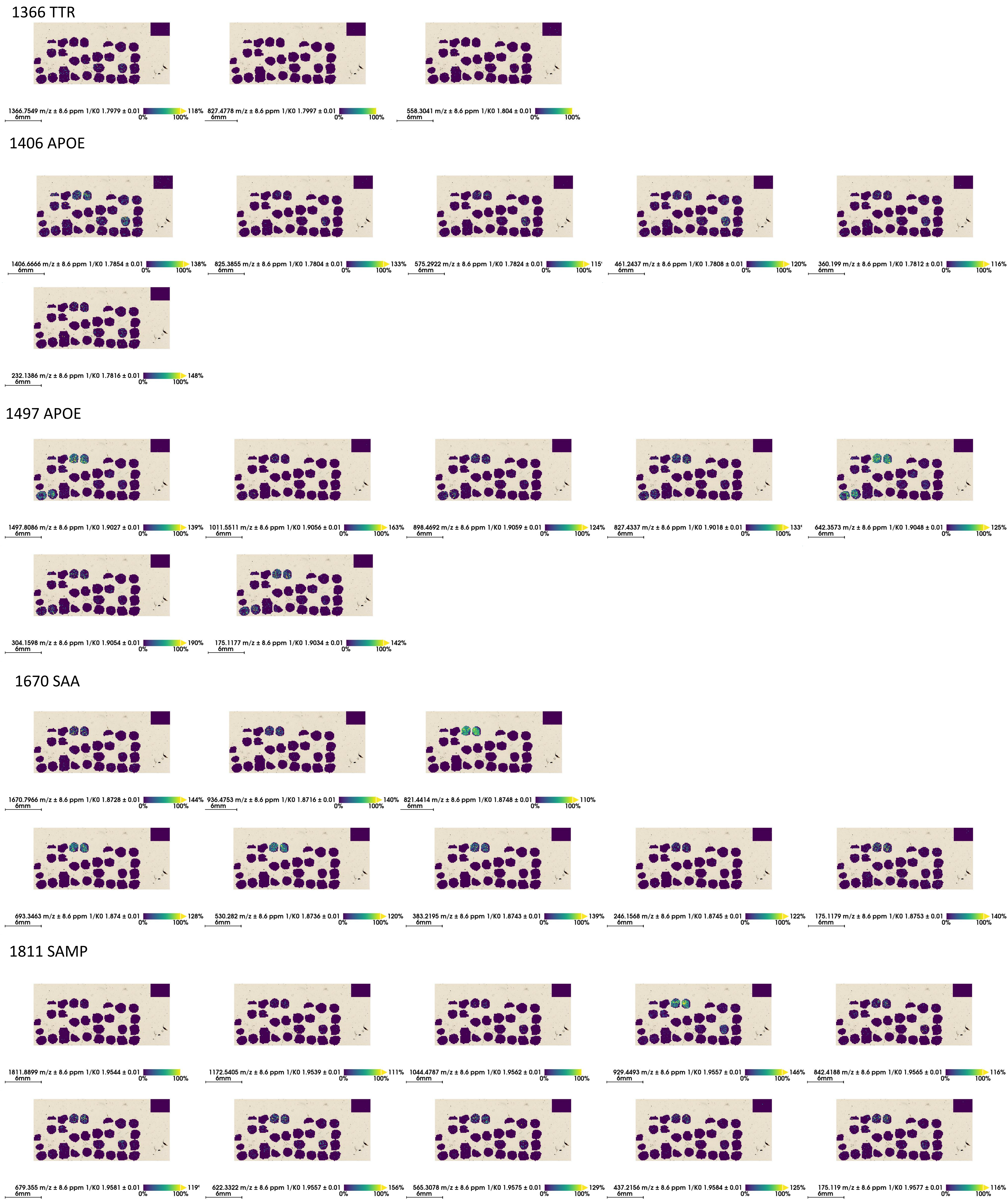


**Figure S4** Continuation.
